# Placental Malaria Induces a Unique Methylation Profile Associated with Fetal Growth Restriction

**DOI:** 10.1101/2024.05.09.593431

**Authors:** Nida Ozarslan, Corina Mong, John Ategeka, Lin Li, Sirirak Buarpung, Joshua F. Robinson, Jimmy Kizza, Abel Kakuru, Moses R. Kamya, Grant Dorsey, Philip J. Rosenthal, Stephanie L. Gaw

## Abstract

**Background:** Fetal growth restriction (FGR) is associated with perinatal death and adverse birth outcomes, as well as long-term complications, including increased childhood morbidity, abnormal neurodevelopment, and cardio-metabolic diseases in adulthood. Placental epigenetic reprogramming associated with FGR may mediate these long-term outcomes. Placental malaria (PM), characterized by sequestration of *Plasmodium falciparum*-infected erythrocytes in placental intervillous space, is the leading global cause of FGR, but its impact on placental epigenetics is unknown. We hypothesized that placental methylomic profiling would reveal common and distinct mechanistic pathways of non-malarial and PM-associated FGR.

**Results:** We analyzed placentas from a US cohort with no malaria exposure (n = 12) and a cohort from eastern Uganda, a region with a high prevalence of malaria (n = 12). From each site, 8 cases of FGR (defined as birth weight <10%ile for gestational age by Intergrowth-21 standard curves) and 4 healthy controls with normal weight were analyzed. PM was diagnosed by placental histopathology. We compared the methylation levels of over 850K CpGs of the placentas using Infinium MethylationEPIC v1 microarray. Non-malarial FGR was associated with 65 differentially methylated CpGs (DMCs), whereas PM-FGR was associated with 133 DMCs, compared to their corresponding controls without FGR. One DMC (cg16389901, located in the promoter region of *BMP4*) was commonly hypomethylated in both groups. We identified 522 DMCs between non-malarial FGR vs. PM-FGR placentas, which was independent of differing geographic location or cellular composition.

**Conclusion:** Placentas with PM-associated FGR have distinct methylation profiles as compared to placentas with non-malarial FGR, suggesting novel epigenetic reprogramming in response to malaria. Larger cohort studies are needed to determine the distinct long-term health outcomes in PM-associated FGR pregnancies.

## Background

Malaria in pregnancy leads to adverse birth outcomes including fetal growth restriction (FGR), low birth weight, preterm birth, and fetal loss. In 2022, 12.7 million pregnant individuals in sub-Saharan Africa were exposed to malaria, potentially resulting in an estimated 914,000 babies being born with low birth weight in the absence of pregnancy-specific chemoprevention Placental malaria (PM), characterized by the sequestration of *Plasmodium falciparum*-infected red blood cells (RBCs) in the placental intervillous spaces, has been independently associated with small for gestational age deliveries and poor neurodevelopmental outcomes (1–4). PM is the leading global cause of FGR. Two mechanisms have been proposed: 1) congestion and inflammation, secondary to the sequestration of infected RBCs in the intervillous spaces during pregnancy, disrupt trophoblast invasion and angiogenesis, decreasing umbilical artery flow (5–7); and 2) vascular dysfunction and chronic inflammation contribute to impaired nutrient exchange at the fetal-maternal interface and imbalances in growth factors (8).

Fetal growth is evaluated by in utero ultrasonographic measurements of fetal biometric measurements (9). FGR is defined as an estimated fetal weight or abdominal circumference less than the 10^th^ percentile for gestational age (10). FGR has important influences on human health. Early onset FGR (<32 weeks gestation) and severe FGR (<3^rd^ percentile) are associated with up to 1.5-fold increased risk of stillbirth and a 2 to 5-fold risk of perinatal death (11). Long-term complications of FGR include increased childhood morbidity and cardiovascular, metabolic, and neuropsychiatric diseases in adulthood (2,4,12–24). In areas with no malaria risk, the etiologies of FGR include maternal comorbidities such as hypertension, autoimmune disease, renal insufficiency as well as placental dysfunction, genetic abnormalities, and environmental exposures (10).

PM and FGR independently impact infant growth and neurodevelopment (25–29). These long-term health impacts may be mediated through epigenetic modifications, including DNA methylation, histone marks, and expression of non-coding RNAs (30–32). For example, placental epigenetic modifications proximal to genes such as IGF2, AHRR, HSD11B2, WNT2, and FOLS1 were found to be associated with FGR (33,34). In contrast, epigenetic studies of human responses to malaria have been largely focused on immune markers of disease susceptibility (35,36). Altered DNA methylation has been demonstrated in monocytes after malaria infection, resulting in age-dependent changes in innate immune and inflammatory responses (37,38). To our knowledge, no studies have examined the epigenetic interactions among FGR and PM-inflicted pregnancies, although existing evidence suggests the potential for shared and divergent mechanisms contributing to altered fetal programming. In this study, we compared methylation profiles in placentas of non-malarial and PM-associated pregnancies, with or without FGR.

## Methods

### Study design and subjects

Placental samples were collected from two independent patient cohorts. Cohort 1 (no malaria) was comprised of 12 placental samples prospectively collected at the University of California, San Francisco, United States (US) from 2018 to 2022 (IRB #10-00505, #20-32077 and #21-33986). Cohort 2 (malaria endemic) was comprised of 12 placental samples prospectively collected from 2021-2022 in Busia, Uganda, as part of a randomized controlled trial of intermittent preventative treatment for malaria in pregnancy (NCT04336189, IRB#19-29105).

Case definitions: All patients were primigravidas and had no evidence of fetal genetic or structural anomalies. FGR cases were identified by estimated fetal weight < 10th percentile for gestational age on all prenatal ultrasounds throughout pregnancy (up to 4 were performed) and birth weight <10th percentile for gestational age by Intergrowth-21 birth weight standards (39). PM-FGR cases were complicated by both FGR and placental malaria (PM) diagnosed by placental histopathology according to Rogerson criteria (40). The presence of *Plasmodium* parasites indicated active PM, whereas malaria pigment (hemozoin) accumulation in the placental tissue was classified as chronic PM (41). Control cases from each cohort were from healthy pregnancies with no pregnancy complications and no evidence of malaria in pregnancy at any time during the study (for Cohort 2). Samples were divided into four groups: 1) FGR (no malarial FGR, n=8), 2) C^FGR^ (healthy controls from the US, n=4), 3) PM-FGR (placental malaria associated FGR, n=8), and 4) C^PM-FGR^ (healthy controls from Uganda, n=4; **Figure 1A**). Detailed case information is provided in **Supplemental Table 1**.

**Figure 1.**
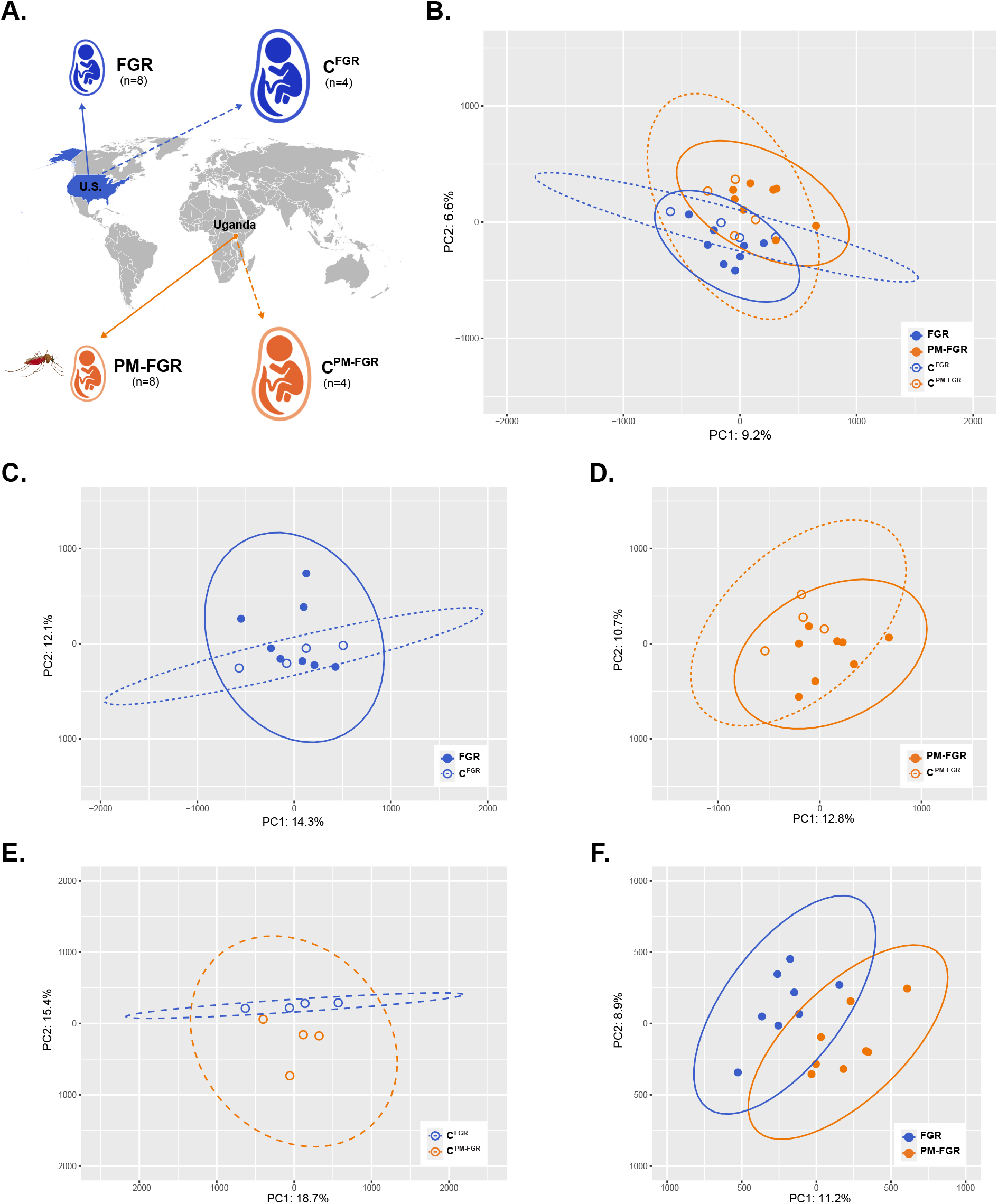
Principal Component Analysis. **A**. Schematic demonstration of study groups and collection sites. **B**. Principal component analysis including all samples. **C**. Principal component analysis of FGR and C^FGR^ samples. **D**. Principal component analysis of PM-FGR and C^PM-FGR^ samples **E**. Principal component analysis of C^FGR^ and C^PM-FGR^ samples **F**. Principal component analysis of FGR and PM-FGR samples.

### Specimen collection

Placental biopsies were collected from the maternal side of the placenta within 1 hour of delivery. Biopsies no larger than 5 × 10 × 10mm were washed with ice-cold PBS three times and placed in RNAlater™ (Qiagen, Germantown, MD). After a minimum incubation of 24 hours in RNAlater at 4°C, biopsies were frozen and stored at -80°C. The same protocol was followed at both sites.

### DNA extraction

DNA was extracted from placental biopsies using a DNeasy Blood & Tissue Kit (Qiagen, Germantown, MD) according to the manufacturer’s standard protocol. Samples were treated with RNase for 2 minutes. Infinite M NanoQuant (Tecan, Morrisville, NC) was used to estimate the purity and concentration of extracted DNA. The A260/A280 ratio for all samples was between 1.75-1.93.

### Infinium MethylationEPIC v1 Assay

We profiled global methylation of DNA aliquots of placental samples. In brief, bisulfate conversion of 1μg DNA/sample was performed using and the EZ DNA Methylation-Lightning Kit (Zymo Research, Irvine, CA). Bisulfite converted DNA was amplified, fragmented, precipitated, resuspended, and hybridized to Infinium Methylation EPIC v1.0 (Illumina, San Diego, CA) arrays following the manufacturer’s protocol. Arrays were washed to remove any unspecific binding, extended and stained primers and imaged using iScan (Illumina, San Diego, CA) platform.

### RNA extraction and qRT-PCR

RNA was extracted from placental biopsies using RNeasy Mini Kit (Qiagen, Germantown, MD) according to the manufacturer’s standard protocol. Infinite M NanoQuant (Tecan, Morrisville, NC) was used to estimate the purity and concentration of extracted RNA. The A260/A280 ratio for all samples was between 2.11-2.19. We converted 500 ng of purified RNA to cDNA using qScript cDNA synthesis kit (Quantabio, Beverly, MA), and performed qRT-PCR using TaqMan primer for *BMP4* (Hs03676628_s1) mixed with TaqMan Universal Master Mix II, no UNG (Life Technologies, Quant Studio 6). Reactions were carried out for 40 cycles. At least 3 technical replicates were analyzed for all comparisons. The expression level of *BMP4* was calculated via the ΔΔCT method. We normalized expression using the mean CT of the housekeeping gene, GAPDH (Hs02786624_g1).

### Bioinformatic analysis

The raw data obtained from the iScan platform was cleaned and preprocessed for data analysis using R (version 4.2.1) and minfi package (version 1.44.0). The quality of the samples was assessed by computing detection p-values for all probes. Low-quality probes that failed at a detection p-value of 0.05 in at least 10% of the samples, probes located on the sex chromosome (X and Y), probes with SNPs at the CpG site, and cross-reactive probes were filtered out prior to quantile normalization and differential methylation analysis. No samples were excluded based on high p-value probes. The total number of probes before filtering was 865,859, whereas total number of probes after filtering was 786,227.

Differential methylation analysis was performed by first computing M values, i.e., the log ratios of methylated vs. unmethylated signal intensities for each probe and limma (version 3.54.2) implemented in the R-package minfi package. Secondly, β-values were calculated, representing the total % methylation level of each CpG site. We performed 4 pairwise comparisons: 1) FGR vs. C^FGR^; 2) PM-FGR vs. C^PM-FGR^; 3) C^FGR^ vs. C^PM-FGR^; 4) FGR vs. PM-FGR. In comparisons among groups, differentially methylated CpGs (DMC) were identified based on an unadjusted *p*-value < 0.0001 and an absolute average difference in methylation (|Δβ|) > 10% (test group vs. control group). Genes proximal to DMCs, were identified using the University of California, Santa Cruz Genome Browser(42). Functional gene ontology (GO) enrichment analysis of genes associated with DMCs was conducted using the Database for Annotation, Visualization and Integrated Discovery (DAVID)(43). For qRT-PCR, we conducted Mann-Whitney test between pairwise comparisons to determine significant differences in expression (p<0.05). Cellular deconvolution analysis was conducted using *planet* (v1.12) package on R (44). Figures were generated using R Studio (v2023.03.0+386), GraphPad Prism (v10.2.3) and BioRender.

## Results

### Characteristics of study participants

To search for associations between FGR types and methylation patterns in samples from the US and Uganda we selected a total of 24 placentas divided into four groups: C^FGR^ (n=4), C^PM-FGR^ (n=4). FGR (n=8), and PM-FGR (n=8). The median and interquartile ranges for gestational age, birth weight and Intergrowth-21 birth weight percentiles and the percentage of female infants were provided in **Table 1**.

**Table 1.** Study group characteristics.

| **Table 1. Study group characteristics.** |  |  |  |  |
| --- | --- | --- | --- | --- |
| **Characteristics** | **FGR**<br>**(n=8)** | **$C^{FGR}$**<br>**(n=4)** | **PM-FGR**<br>**(n=8)** | **$C^{PM-FGR}$**<br>**(n=4)** |
| Gestational age | 37.5<br>(36.7-38.2) | 39.2<br>(39.1-39.5) | 39.8<br>(39.1-41.2) | 39.2<br>(38.3-40.1) |
| Birth weight (gr) | 2193<br>(2004-2355) | 3565<br>(3048-3768) | 2600<br>(2510-2843) | 2915<br>(2885-3470) |
| Intergrowth-21 birth weight percentile | 2.9<br>(1.1-5.5) | 80.1<br>(30.1-87.7) | 3.9<br>(3.2-5.7) | 29.4<br>(21.8-76.1) |
| Infant sex (female) | 62.5% | 25% | 50% | 75% |

The median and interquartile ranges were provided for gestational age, birth weight and Intergrowth-21 birth weight percentiles. Percentage of female infants were provided for infant sex.

### Dimensional Reduction Analysis

We profiled the methylome of our selected placentas using Illumina Infinium MethylationEPIC Arrays v1. After applying quality control measures, we compared methylation patterns of 786,227 CpG sites among FGR, PM-FGR, and respective controls, C^FGR^ and C^PM-FGR^, from the two unique geographic locations. Unsupervised principal component analysis using M-values of all CpG sites suggested no overall distinct separation among the four groups, suggesting general similarities in global methylation profiles independent of FGR or PM-FGR status or collection site (**Figure 1B-E**). In pairwise comparisons between groups, separation was most prominent between FGR and PM-FGR groups (**Figure 1F**).

### Identification of Differentially Methylated CpGs

To better characterize differences in methylation levels with non-malarial FGR and PM-associated FGR we investigated the differentially methylated CpGs using an unadjusted p value cutoff of p<0.0001 and a secondary cutoff of at least 10% absolute difference in mean methylation (|Δβ| > 0.10). To confirm the power of the significance cutoff in terms of differentiating the groups, unsupervised hierarchical clustering was performed using β-values of the respective DMCs. As expected, our samples clustered based on study group (**Supplemental Figure 1, Supplemental Data 1**).

### Differentially Methylated CpGs with FGR

We first investigated the influence of non-malarial FGR on placental methylation profiles in comparison with their respective geographic controls. We identified 65 DMCs, that were distributed across all autosomal chromosomes except for chromosomes 19 and 21 (**Figure 2A**). Of the 65 DMCs, 48 were hypomethylated and 17 were hypermethylated with non-malarial FGR vs. C^FGR^ (**Figure 2B**). While most of the DMCs were located at the body and intergenic regions, 14 hypomethylated CpGs and 6 hypermethylated CpGs were located in the promoter or 1^st^ exon region (**Figure 2C)**. Functional enrichment analysis through DAVID revealed that the genes associated with the 14 hypomethylated CpGs located in the promoter or 1^st^ exon region were involved in transport, cellular response to stress, nephron development and localization within membrane; in contrast, the 6 hypermethylated CpGs were associated with genes involved in angiogenesis.

**Figure 2.**
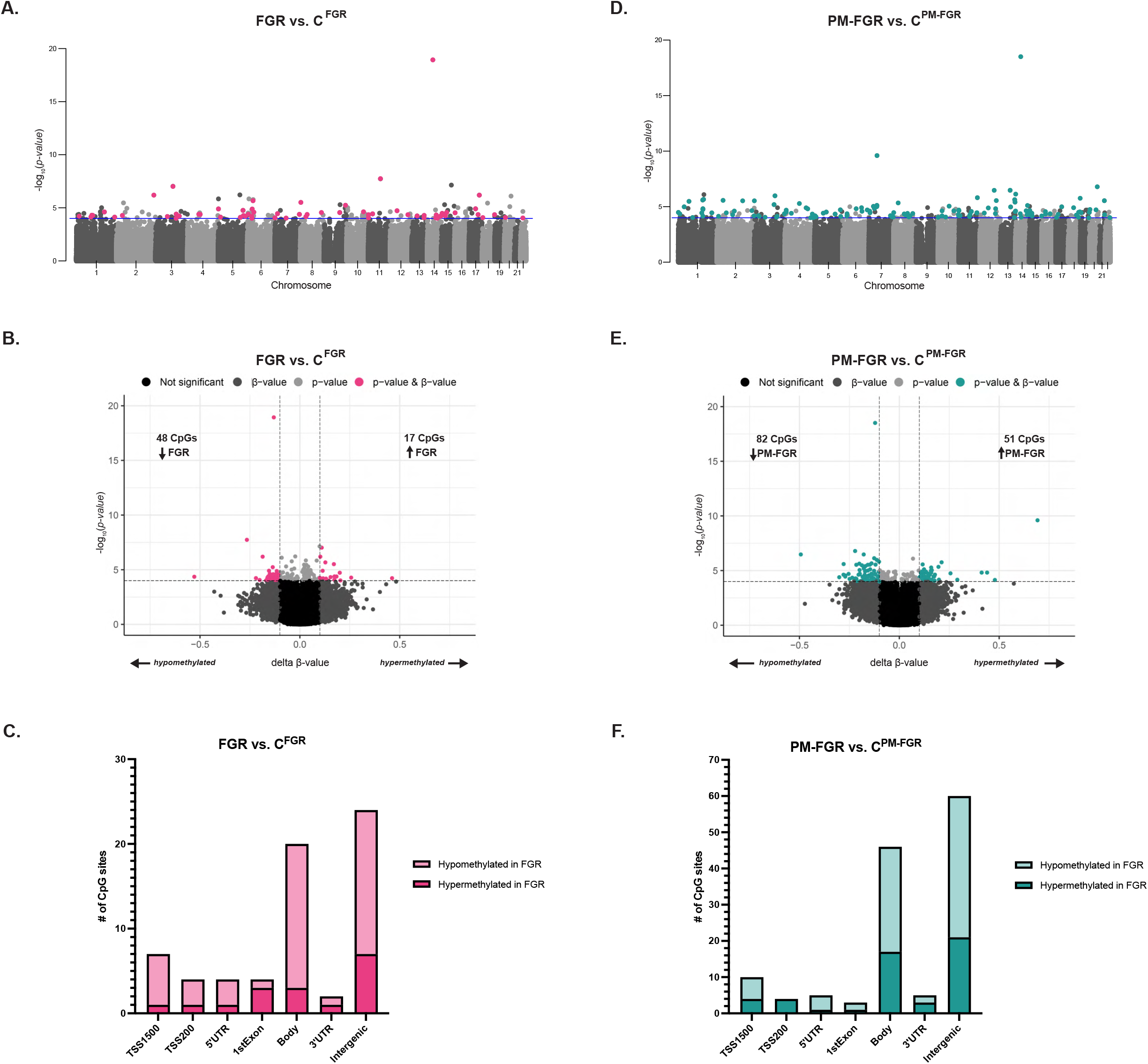
Differentially methylated CpGs (DMCs) observed with non-malarial FGR and placental malaria-associated FGR and their respective controls. **A**. Manhattan plot of 65 DMCs (p<0.0001 and |Δβ| > 0.1) between FGR vs. C^FGR^ demonstrated by pink circles. **B**. Volcano plot displaying 48 hypomethylated and 17 were hypermethylated CpGs with FGR vs. C^FGR^ indicated by pink circles. **C**. Bar graph demonstrating the distribution of hypermethylated (dark pink) and hypomethylated (light pink) DMCs with FGR vs. C^FGR^ among CpG positions. **D**. Manhattan plot of 133 DMCs (p<0.0001 and |Δβ| > 0.1) between PM-FGR vs. C^PM-FGR^ demonstrated by teal circles. **E**. Volcano plot displaying 82 hypomethylated and 51 were hypermethylated CpGs with PM-FGR vs. C^PM-FGR^ indicated by teal circles. **F**. Bar graph demonstrating the distribution of hypermethylated (dark teal) and hypomethylated (light teal) DMCs with PM-FGR vs. C^PM-FGR^ among CpG positions.

### Differentially Methylated CpGs with PM-FGR

Next, we investigated the DMCs between PM-FGR vs. C^PM-FGR^, we identified 133 CpGs that were distributed across all autosomal chromosomes with the exception of chromosome 9 (**Figure 2D**). Of the 133 DMCs, 82 were hypomethylated and 51 were hypermethylated with non-malarial FGR vs. C^FGR^ (**Figure 2E**). While most of the DMCs were located at the body and intergenic regions, 12 hypomethylated CpGs and 10 hypermethylated CpGs were located in the promoter or 1^st^ exon region (**Figure 2F)**. Genes associated with the 12 hypomethylated CpGs located in the promoter or 1^st^ exon region were involved in signaling; the 6 hypermethylated CpGs were associated with genes involved in cell differentiation, tissue development and transport across the membrane.

### Common changes observed with non-malarial FGR and PM-FGR

To identify the epigenetic modulations observed with both non-malarial and PM-FGR, we investigated the overlapping DMCs between FGR vs. C^FGR^ and PM-FGR vs. C^PM-FGR^ comparisons. We identified only a single CpG, cg16389901, that was commonly hypomethylated with both FGR and PM-FGR when compared to their respective controls (**Figure 3A**). In comparing trends in methylation patterns among DMCs identified in either disease group, we observed distinct patterns between non-malarial FGR and PM-FGR (**Figure 3B-3C**). The commonly hypomethylated CpG site cg16389901 is in the promoter region of two splice variants of *BMP4*, a cell growth-related gene known to participate in trophoblast differentiation within the placenta (45) (**Figure 3D**). Interestingly, DNA methylation levels of other CpGs associated with *BMP4* did not show a difference between FGR vs. C^FGR^ or PM-FGR vs. C^PM-FGR^ comparisons, suggesting a specific association with FGR. To understand the impact of differential DNA methylation on the mRNA level we conducted qRT-PCR targeting *BMP4*. The mRNA expression levels of *BMP4* did not show any significant difference between the four study groups (**Figure 3E**).

**Figure 3.**
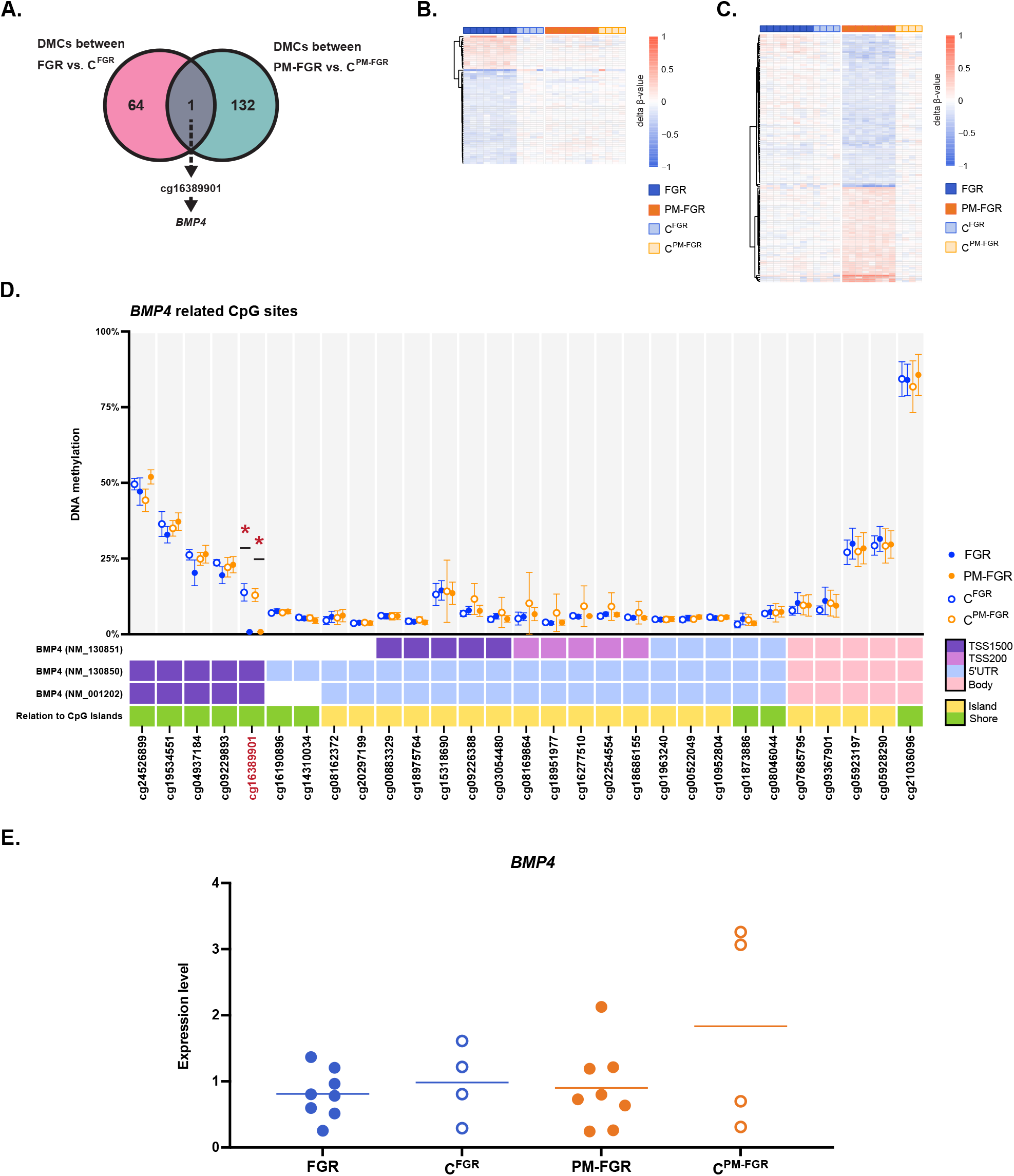
Commonly differentially methylated CpG site with non-malarial FGR and placental malaria-associated FGR. **A**. Venn diagram demonstrating the sole overlapping differentially methylated CpG site, cg16389901, that was commonly hypomethylated with both non-malarial and placental malaria-associated FGR when compared to their controls. **B**. Normalized methylation levels of all samples based on their geographically matched control group average of 65 DMCs identified between FGR vs. C^FGR^ **C**. Normalized methylation levels of all samples based on their geographically matched control group average of 133 DMCs identified between PM-FGR vs. C^PM-FGR^ **D**. Average DNA methylation of CpG sites associated with *BMP4*, including cg16389901 (marked in red), for all four groups. Red asterisk indicates significance. **E**. Gene expression levels of *BMP4* measured by qRT-PCR.

### Differentially methylated CpGs between FGR vs. PM-FGR

We next compared the methylation profiles between FGR and PM-FGR. To account for baseline epigenetic differences based on the distinct geographical locations of the two cohorts, we compared the DMCs between the two control groups (C^FGR^ vs. C^PM-FGR^), showing 183 CpGs that were differentially methylated between the two study sites. Of these 183 DMCs, 7 CpG sites were also identified when comparing FGR vs. PM-FGR. The exclusion of these 7 CpGs as potentially affected by population-based differences resulted in 522 CpGs that were associated with PM status (**Figure 4A**). The number of DMCs identified between FGR vs. PM-FGR was more than 8-fold than identified between FGR vs. C^FGR^ and 3-fold between PM-FGR vs. C^PM-FGR^ comparisons, highlighting the distinct changes observed with PM. The 522 DMCs were distributed across all autosomal chromosomes (**Figure 4B**). Of the 522 DMCs, 143 were hypermethylated in non-malarial FGR placentas and 379 were hypermethylated in PM-FGR placentas (**Figure 4C**). While most of the DMCs were located at the body and intergenic regions, 30 hypermethylated CpGs with FGR and 106 CpGs hypermethylated with PM-FGR were located in the promoter or 1^st^ exon region (**Figure 4D)**. Genes associated with the 30 hypermethylated CpGs with FGR located in the promoter or 1^st^ exon region were involved in actin cytoskeleton organization, whereas the 106 hypermethylated CpGs with PM-FGR were associated with genes involved in multicellular organism development, regulation of metabolic processes, nervous system process, cell differentiation and response to growth factor. Particularly, 3 CpG sites in the promoter region of *ARPC1B* and 4 CpG sites in the promoter region of *ZSCAN23* were hypermethylated with FGR. On the other hand, 7 CpG sites in the promoter region of *FAM124B* and 4 CpG sites in the promoter region of *SPG7* were hypermethylated with PM-FGR.

**Figure 4.**
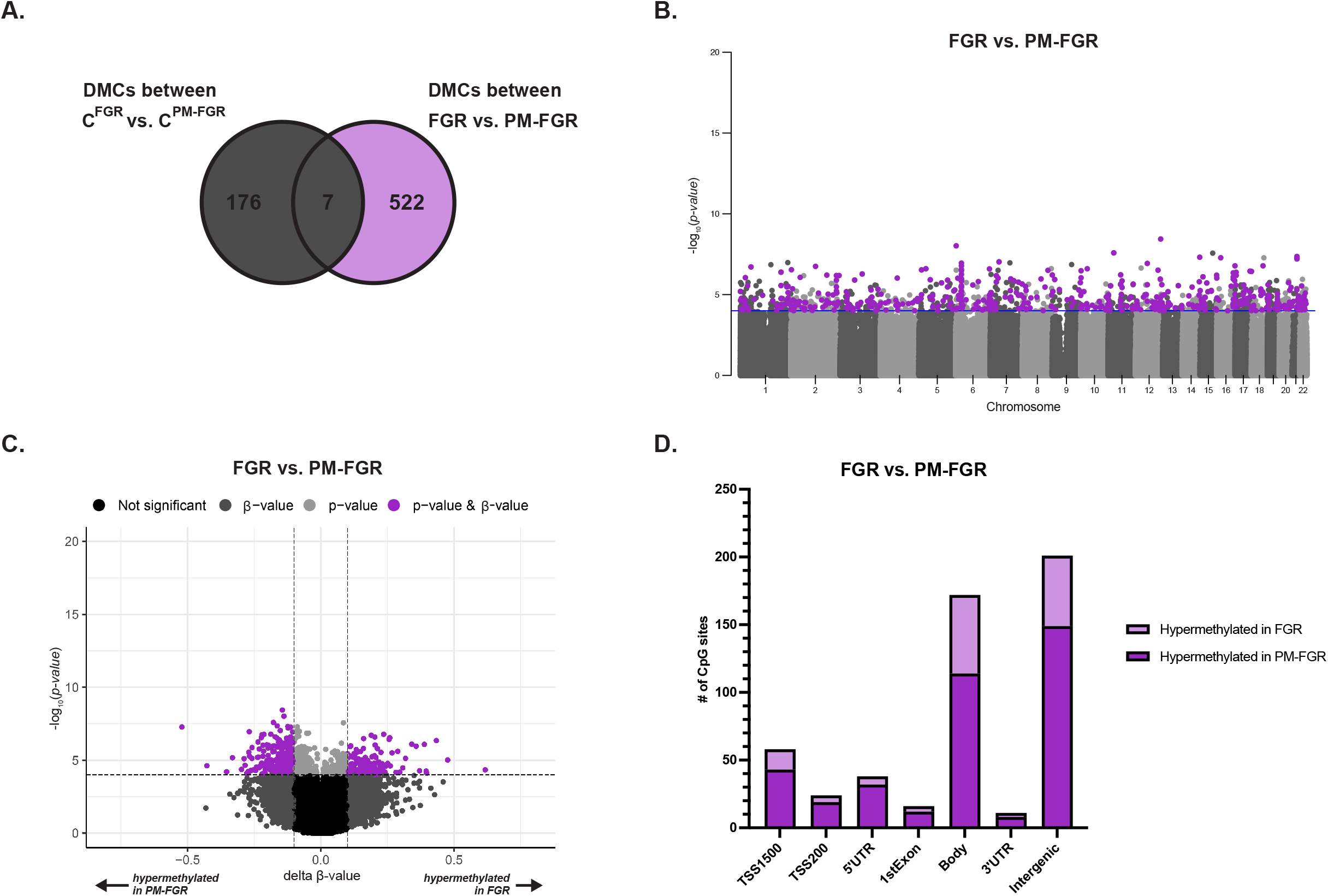
Differentially methylated CpGs (DMCs) observed between non-malarial FGR vs. placental malaria-associated FGR. **A**. Venn diagram demonstrating the 522 DMCs (p<0.0001 and |Δβ| > 0.1) between FGR vs. PM-FGR after subtracting geographical location-driven baseline differences **B**. Manhattan plot of 522 DMCs between FGR vs. PM-FGR demonstrated by purple circles. **C**. Volcano plot displaying 143 hypermethylated CpGs in FGR and 379 CpGs hypermethylated in PM-FGR indicated by purple circles. **D**. Bar graph demonstrating the distribution of hypermethylated CpGs with FGR (light purple) and hypermethylated with PM-FGR (dark purple) among CpG positions.

### Identifying the cellular composition of all placental samples

Our study used DNA extracted from homogenized placental tissue. To understand the cellular composition of each sample, we performed cellular deconvolution analysis based on reference methylation data of term placental cells generated by Yuan et al. (44) (**Figure 5A**). The robust partial correlations (RPC) method was used to identify the percentages of syncytiotrophoblasts, trophoblasts, endothelial cells, stromal cells, Hofbauer cells and nucleated red blood cells (nRBC) based on β-values of the most distinguishing CpG sites among these cell types. As expected, our placental samples were predicted based to be enriched for syncytiotrophoblasts, constituting an average of 69.5% of all cells. The next most prevalent cell types were stromal cells (13.9%), endothelial cells (12.1%), and trophoblasts (3.8%). The average abundance of Hofbauer cells and nRBC in total was less than 0.6%. We performed unsupervised hierarchical clustering to investigate whether the cellular composition of samples differed between the four study groups (**Figure 5B**). Overall, the cellular composition of samples did not show any clear separation between groups (**Figure 5B, Supplemental Figure 2**). However, hierarchical clustering of only FGR and PM-FGR samples demonstrated a more prominent clustering between the two groups (**Figure 5C**). This separation was primarily associated with a significant increase in the percentage of syncytiotrophoblasts in PM-FGR compared to FGR samples (74.4% vs. 65.5%, respectively; p<0.002). Only a single CpG site overlapped with the 522 identified CpGs used for cellular deconvolution analysis, indicating that DMCs identified between FGR vs. PM-FGR were unlikely to be driven by differences in cellular composition (**Figure 5D**).

**Figure 5.**
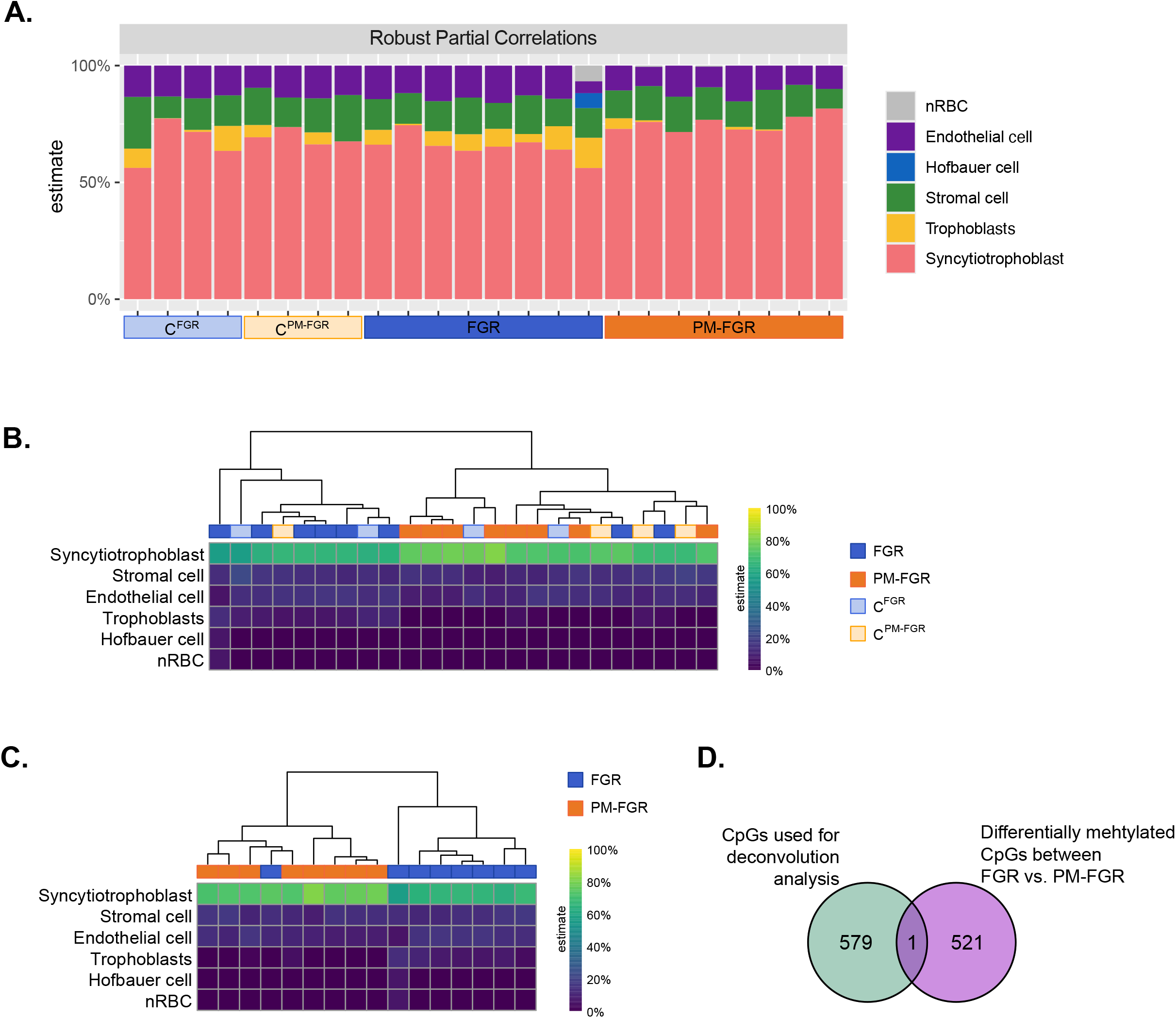
Cellular deconvolution analysis based on overall methylation levels. **A**. Cell composition of all samples estimated using robust partial correlations analysis through planet package in R. **B**. Hierarchical clustering of all samples based on cellular deconvolution results. **C**. Hierarchical clustering of FGR and PM-FGR samples based on cellular deconvolution analysis. **D**. Venn diagram demonstrating the overlap between CpGs used for cellular deconvolution analysis and differentially methylated CpGs between FGR vs. PM-FGR.

## Discussion

To our knowledge, this is the first study to investigate methylation profiles in PM. We found that placentas from non-malarial FGR and placental malaria-associated FGR cases exhibited highly divergent methylation profiles. Interestingly, there was only a single CpG site, associated with the promoter region of *BMP4*, that was commonly hypomethylated in placentas associated with both non-malarial FGR and PM-FGR. In PM-FGR, hypermethylated CpGs were associated with genes encoding proteins involved in developmental processes, whereas hypermethylated CpGs in FGR were associated with genes encoding proteins involved in metabolic processes. Cellular deconvolution demonstrated differences in the relative abundances of placental cell types between placentas associated with non-malarial and PM-FGR; however, these differences did not explain the distinct methylation profiles of these groups. Overall, our results suggest novel epigenetic modifications in response to PM-associated FGR compared to those seen with non-malarial FGR.

Previous studies investigating placental epigenetic changes related to FGR in populations without risk of malaria identified various differentially methylated CpG sites in genes encoding proteins involved with fatty acid oxidation, transcriptional regulation, immune responses, cell adhesion, metabolism, and cell growth (33,46–49). Similar to our study, one study from Korea investigating the methylation levels of placenta and cord blood samples with FGR described changes in methylation profiles of genes encoding proteins involved with metabolism and developmental pathways (48). While various studies investigated methylation profiles of placenta, maternal blood or cord blood samples from pregnancies complicated with fetal growth restriction, there was a lack of overlap of differentially methylated CpG sites as well as genes related to those DMCs across these studies (33,34,46,47,49–52). Our results revealed unique differentially methylated CpG sites and related genes compared to the previous FGR related studies. Differences in results between studies may have been due to diverse mechanisms of epigenetic modulation other than DNA methylation, such as histone modifications or non-coding RNA, as well as differences in study populations.

In analyses of placentas, both non-malarial FGR and PM-FGR were associated with moderate numbers of differentially methylated CpGs compared to their corresponding controls. Interestingly, a single CpG site was commonly hypomethylated with both non-malarial FGR and PM-FGR—cg16389901. This site is located 200-1500 bases upstream of the transcription start site of the *BMP4* gene. *BMP4* is a well-studied growth-associated protein that also plays an important role in trophoblast differentiation (45,53). The Korean study also described hypermethylation of genes encoding proteins involved in the negative regulation of the BMP signaling pathway, which aligns with our results (48). Hypomethylation of *BMP4*, resulting in increased expression, may be a common compensatory mechanism in the placenta to counteract other functional changes that lead to inhibited fetal growth. In our placental samples, mRNA levels of *BMP4* did not show any significant difference between our study groups. This could suggest that the hypomethylation of *BMP4* with both non-malarial FGR and PM-FGR is a compensatory response to normalize the mRNA levels in the setting of other mechanisms that may result in reduced expression of this protein (e.g. histone modifications, posttranslational modifications)

Hypermethylated CpG sites in placentas from non-malarial FGR compared to PM-associated FGR included CpGs located at the promoter regions of *ARPC1B* and *ZSCAN23*. Homozygous mutations in the *ARPC1B* gene have been associated with disruptions in the immune system (54). Promoter region hypermethylation of *ZSCAN23* has been associated with the acceleration of pancreatic cancer growth (55). Conversely, hypermethylated CpG sites in placentas from PM-associated compared to non-malarial FGR included CpGs located at the promoter regions of *FAM124B* and *SPG7. FAM124B* has been proposed as an important protein involved in neurodevelopmental disorders, while SpG7 mutations have been linked to spastic paraplegia and cerebellar ataxia (56,57). Further follow-up studies on offspring are needed to investigate long-term outcomes associated with non-malarial FGR vs. PM-associated FGR and to understand whether these outcomes are mediated by epigenetic mechanisms.

Cellular deconvolution analysis revealed dominance of syncytiotrophoblasts, which were present in significantly higher proportions in PM-associated FGR samples. Differences in the proportion of syncytiotrophoblasts may be associated with a classic pathological finding of syncytial knot formation commonly observed with PM, which is thought to be a response to inflammatory damage associated with infected RBCs in the intervillous spaces (58). Despite the differences in cellular composition, the differentially methylated CpGs were distinct from those associated with specific cell types. In contrast, a study that conducted the same deconvolution analysis on placentas from monochorionic twins with selective FGR found decreased proportions of stromal cells in the twin with FGR compared to the normally grown twin (46). This suggests that placental cellular composition may change in different etiologies of FGR and warrants further study.

Our study has some limitations. First, we had a small sample size, and larger studies are needed to confirm our findings. Second, due to the homogenization of placental tissue, we were unable to distinguish between the origins of different cell types, particularly maternal and fetal immune cells. Third, due to the high prevalence of malaria in pregnancy at our study site in Uganda, we did not identify any FGR cases from Uganda without evidence of malaria infection during pregnancy; these cases would provide a more direct comparison between non-malarial and PM-associated FGR in an African population. Finally, we focused on methylation profiles detected by a targeted methylation array, rather than whole genome bisulfide sequencing, which may identify additional differentially methylated CpGs. Further studies investigating other types of epigenetic mechanisms such as histone modifications or assessment of open chromatin regions would provide deeper insight into epigenetic changes occurring with non-malarial and PM-associated FGR. Additional transcriptomic and proteomic analysis of similar cohorts would also reveal insight into downstream effects of epigenetic modifications.

This is the first study, to our knowledge, comparing the methylation profiles of malarial and non-malarial FGR in the placenta. Other strengths of our study include detailed characterizations of the pregnancies, allowing for the selection of appropriate controls for each population. We only included primigravid patients to avoid confounding from gravidity-induced differences in methylation profiles. Finally, placental biopsies from both sites were collected and processed with the same protocol, allowing for comparison between the different patient populations.

The significant differences that we found in DMCs in non-malarial and PM-associated FGR, even after controlling for geographic location, suggest distinct pathways of epigenetic reprogramming in the setting of FGR with or without PM. The independent and combined impact of placental malaria and placental epigenetics may have potential implications for long-term health in the offspring.

## Declarations

### Ethics approval and consent to participate

This study was approved by the University of California, San Francisco and Makerere University. Written informed consent was obtained from all participants.

### Availability of data and materials

The dataset generated during the study will be available in the GEO repository (waiting for accession number).

### Competing interests

The authors declare that they have no competing interests.

### Funding

This study was supported by the National Institutes of Health (NIAID K08AI141728 to SLG and U01AI141308 to GD and PJR), and the Bill and Melinda Gates Foundation (INV-017035 to SLG).

### Authors’ contributions

Conceptualization: NO and SLG Methodology: NO and SLG Investigation: NO, CM, JA, LL, SB, JFR Funding acquisition: MRK, GD, PJR, SLG Project administration: AK, JK Supervision: AK, SLG Writing – original draft: NO and SLG Writing – review & editing: All Authors

#### Acknowledgments

The authors thank all study participants. We also thank Keith Boohrer, Xiaojing Yang and Kai Chang from the Zymo Research Corporation for their technical support.

## Supplementary Documents

**Supplementary Table 1. Patient characteristics**. cHTN; chronic hypertension, gHTN; gestational hypertension, gDM; gestational DM, PreE; preeclampsia, SIPE; superimposed preeclampsia.

**Supplementary Figure 1. Hierarchical clustering of samples based on methylation levels of differentially methylated CpGs (DMCs**). A. Methylation level of 65 DMCs between FGR vs. C^FGR^ **B**. Methylation level of 133 DMCs between PM-FGR vs. C^PM-FGR^ **C**. Methylation level of 183 DMCs between C^FGR^ vs. C^PM-FGR^ **D**. Methylation level of 522 DMCs between FGR vs. PM-FGR

**Supplementary Figure 2. Hierarchical clustering of samples based on cellular composition. A**. Hierarchical clustering of FGR and C^FGR^ samples based on cellular composition. **B**. Hierarchical clustering of PM-FGR and C^PM-FGR^ samples based on cellular composition.

**Supplementary Data 1. List of differentially methylated CpG sites**.

## Notes

### Competing Interest Statement

The authors have declared no competing interest.

### Summary of Updates

Figure 2, Manhattan Plot (panel A&D) has been updated, new bar graphs describing the distribution of positions of DMCs (panel C&F) have been added. Figure 3, heatmap of DMCs (panel B&C) and qRT-PCR results for BMP4 (panel E) have been added. Figure 4, Manhattan Plot (panel B) has been updated, new bar graph describing the distribution of positions of DMCs (panel C) has been added. Former Table 1 has been moved to supplementary documents and new Table 1 has been added. Abstract, introduction, methods, results and discussion sections have been updated accordingly.

## References

1. Garrison A, Boivin MJ, Fiévet N, Zoumenou R, Alao JM, Massougbodji A, et al. The Effects of Malaria in Pregnancy on Neurocognitive Development in Children at 1 and 6 Years of Age in Benin: A Prospective Mother-Child Cohort. Clin Infect Dis Off Publ Infect Dis Soc Am. 2022 Mar 9;74(5):766–75.

2. Bangirana P, Conroy AL, Opoka RO, Semrud-Clikeman M, Jang JH, Apayi C, et al. Effect of Malaria and Malaria Chemoprevention Regimens in Pregnancy and Childhood on Neurodevelopmental and Behavioral Outcomes in Children at 12, 24, and 36 Months: A Randomized Clinical Trial. Clin Infect Dis. 2023 Feb 18;76(4):600–8.

3. Conroy AL, Bangirana P, Muhindo MK, Kakuru A, Jagannathan P, Opoka RO, et al. Case Report: Birth Outcome and Neurodevelopment in Placental Malaria Discordant Twins. Am J Trop Med Hyg. 2019 Mar;100(3):552–5.

4. Bangirana P, Opoka RO, Boivin MJ, Idro R, Hodges JS, Romero RA, et al. Severe Malarial Anemia is Associated With Long-term Neurocognitive Impairment. Clin Infect Dis. 2014 Aug 1;59(3):336–44.

5. Moore KA, Simpson JA, Wiladphaingern J, Min AM, Pimanpanarak M, Paw MK, et al. Influence of the number and timing of malaria episodes during pregnancy on prematurity and small-for-gestationalage in an area of low transmission. BMC Med. 2017 Dec;15(1):117.

6. Chua CLL, Khoo SKM, Ong JLE, Ramireddi GK, Yeo TW, Teo A. Malaria in Pregnancy: From Placental Infection to Its Abnormal Development and Damage. Front Microbiol. 2021 Nov 11;12:777343.

7. Dorman EK, Shulman CE, Kingdom J, Bulmer JN, Mwendwa J, Peshu N, et al. Impaired uteroplacental blood flow in pregnancies complicated by falciparum malaria: Uterine artery Doppler and malaria. Ultrasound Obstet Gynecol. 2002 Feb;19(2):165–70.

8. Chandrasiri UP, Chua CLL, Umbers AJ, Chaluluka E, Glazier JD, Rogerson SJ, et al. Insight Into the Pathogenesis of Fetal Growth Restriction in Placental Malaria: Decreased Placental Glucose Transporter Isoform 1 Expression. J Infect Dis. 2014 May 15;209(10):1663–7.

9. Hadlock FP, Harrist RB, Martinez-Poyer J. In utero analysis of fetal growth: a sonographic weight standard. Radiology. 1991 Oct;181(1):129–33.

10. ACOG Practice Bulletin No. 204: Fetal Growth Restriction. Obstet Gynecol. 2019 Feb;133(2):e97– 109.

11. Madden JV, Flatley CJ, Kumar S. Term small-for-gestational-age infants from low-risk women are at significantly greater risk of adverse neonatal outcomes. Am J Obstet Gynecol. 2018 May;218(5):525.e1-525.e9.

12. Weckman AM, Conroy AL, Madanitsa M, Gnaneswaran B, McDonald CR, Kalilani-Phiri L, et al. Neurocognitive outcomes in Malawian children exposed to malaria during pregnancy: An observational birth cohort study. PLoS Med. 2021 Sep;18(9):e1003701.

13. Colella M, Frérot A, Novais ARB, Baud O. Neonatal and Long-Term Consequences of Fetal Growth Restriction. Curr Pediatr Rev. 2018 Dec 21;14(4):212–8.

14. Lesseur C, Paquette AG, Marsit CJ. Epigenetic Regulation of Infant Neurobehavioral Outcomes. Med Epigenetics. 2014 Jun 11;2(2):71–9.

15. Mericq V, Martinez-Aguayo A, Uauy R, Iñiguez G, Van Der Steen M, Hokken-Koelega A. Long-term metabolic risk among children born premature or small for gestational age. Nat Rev Endocrinol. 2017 Jan;13(1):50–62.

16. D’Agostin M, Di Sipio Morgia C, Vento G, Nobile S. Long-term implications of fetal growth restriction. World J Clin Cases. 2023 May 6;11(13):2855–63.

17. Chan PYL, Morris JM, Leslie GI, Kelly PJ, Gallery EDM. The Long-Term Effects of Prematurity and Intrauterine Growth Restriction on Cardiovascular, Renal, and Metabolic Function. Int J Pediatr. 2010;2010:1–10.

18. Leon DA, Lithell HO, Vagero D, Koupilova I, Mohsen R, Berglund L, et al. Reduced fetal growth rate and increased risk of death from ischaemic heart disease: cohort study of 15 000 Swedish men and women born 1915-29. BMJ. 1998 Jul 25;317(7153):241–5.

19. Martyn CN, Barker DJ, Jespersen S, Greenwald S, Osmond C, Berry C. Growth in utero, adult blood pressure, and arterial compliance. Heart. 1995 Feb 1;73(2):116–21.

20. Liefke J, Heijl C, Steding-Ehrenborg K, Morsing E, Arheden H, Ley D, et al. Fetal growth restriction followed by very preterm birth is associated with smaller kidneys but preserved kidney function in adolescence. Pediatr Nephrol Berl Ger. 2023 Jun;38(6):1855–66.

21. Senra JC, Yoshizaki CT, Doro GF, Ruano R, Gibelli MABC, Rodrigues AS, et al. Kidney impairment in fetal growth restriction: three-dimensional evaluation of volume and vascularization. Prenat Diagn. 2020 Oct;40(11):1408–17.

22. Arigliani M, Stocco C, Valentini E, De Pieri C, Castriotta L, Ferrari ME, et al. Lung function between 8 and 15 years of age in very preterm infants with fetal growth restriction. Pediatr Res. 2021 Sep;90(3):657–63.

23. Nikolajev K, Heinonen K, Hakulinen A, Länsimies E. Effects of intrauterine growth retardation and prematurity on spirometric flow values and lung volumes at school age in twin pairs. Pediatr Pulmonol. 1998 Jun;25(6):367–70.

24. Grunnet LG, Bygbjerg IC, Mutabingwa TK, Lajeunesse-Trempe F, Nielsen J, Schmiegelow C, et al. Influence of placental and peripheral malaria exposure in fetal life on cardiometabolic traits in adult offspring. BMJ Open Diabetes Res Care. 2022 Apr;10(2):e002639.

25. Korzeniewski SJ, Allred EN, Joseph RM, Heeren T, Kuban KCK, O’Shea TM, et al. Neurodevelopment at Age 10 Years of Children Born & lt;28 Weeks With Fetal Growth Restriction. Pediatrics. 2017 Nov 1;140(5):e20170697.

26. Leitner Y, Fattal-Valevski A, Geva R, Eshel R, Toledano-Alhadef H, Rotstein M, et al. Neurodevelopmental outcome of children with intrauterine growth retardation: a longitudinal, 10-year prospective study. J Child Neurol. 2007 May;22(5):580–7.

27. Murray E, Fernandes M, Fazel M, Kennedy S, Villar J, Stein A. Differential effect of intrauterine growth restriction on childhood neurodevelopment: a systematic review. BJOG Int J Obstet Gynaecol. 2015 Jul;122(8):1062–72.

28. Elgueta D, Murgas P, Riquelme E, Yang G, Cancino GI. Consequences of Viral Infection and Cytokine Production During Pregnancy on Brain Development in Offspring. Front Immunol. 2022 Apr 7;13:816619.

29. Sacchi C, Marino C, Nosarti C, Vieno A, Visentin S, Simonelli A. Association of Intrauterine Growth Restriction and Small for Gestational Age Status With Childhood Cognitive Outcomes: A Systematic Review and Meta-analysis. JAMA Pediatr. 2020 Aug 1;174(8):772.

30. Lapehn S, Paquette AG. The Placental Epigenome as a Molecular Link Between Prenatal Exposures and Fetal Health Outcomes Through the DOHaD Hypothesis. Curr Environ Health Rep. 2022 Apr 29;9(3):490–501.

31. Nawathe AR, Christian M, Kim SH, Johnson M, Savvidou MD, Terzidou V. Insulin-like growth factor axis in pregnancies affected by fetal growth disorders. Clin Epigenetics. 2016 Dec;8(1):11.

32. Lee MH, Jeon YJ, Lee SM, Park MH, Jung SC, Kim YJ. Placental gene expression is related to glucose metabolism and fetal cord blood levels of insulin and insulin-like growth factors in intrauterine growth restriction. Early Hum Dev. 2010 Jan;86(1):45–50.

33. Xiao X, Zhao Y, Jin R, Chen J, Wang X, Baccarelli A, et al. Fetal growth restriction and methylation of growth-related genes in the placenta. Epigenomics. 2016 Jan;8(1):33–42.

34. Tekola-Ayele F, Zeng X, Ouidir M, Workalemahu T, Zhang C, Delahaye F, et al. DNA methylation loci in placenta associated with birthweight and expression of genes relevant for early development and adult diseases. Clin Epigenetics. 2020 Dec;12(1):78.

35. Arama C, Quin JE, Kouriba B, Östlund Farrants AK, Troye-Blomberg M, Doumbo OK. Epigenetics and Malaria Susceptibility/Protection: A Missing Piece of the Puzzle. Front Immunol. 2018 Aug 3;9:1733.

36. Goldberg AD, Allis CD, Bernstein E. Epigenetics: A Landscape Takes Shape. Cell. 2007 Feb;128(4):635–8.

37. Dobbs KR, Embury P, Koech E, Ogolla S, Munga S, Kazura JW, et al. Age-related differences in monocyte DNA methylation and immune function in healthy Kenyan adults and children. Immun Ageing. 2021 Dec;18(1):11.

38. Romero DVL, Balendran T, Hasang W, Rogerson SJ, Aitken EH, Achuthan AA. Epigenetic and transcriptional regulation of cytokine production by Plasmodium falciparum-exposed monocytes. Sci Rep. 2024 Feb 5;14(1):2949.

39. Papageorghiou AT, Ohuma EO, Altman DG, Todros T, Ismail LC, Lambert A, et al. International standards for fetal growth based on serial ultrasound measurements: the Fetal Growth Longitudinal Study of the INTERGROWTH-21st Project. The Lancet. 2014 Sep;384(9946):869–79.

40. Rogerson SJ, Hviid L, Duffy PE, Leke RF, Taylor DW. Malaria in pregnancy: pathogenesis and immunity. Lancet Infect Dis. 2007 Feb;7(2):105–17.

41. Zakama AK, Ozarslan N, Gaw SL. Placental Malaria. Curr Trop Med Rep. 2020 Dec;7(4):162–71.

42. Nassar LR, Barber GP, Benet-Pagès A, Casper J, Clawson H, Diekhans M, et al. The UCSC Genome Browser database: 2023 update. Nucleic Acids Res. 2023 Jan 6;51(D1):D1188–95.

43. Sherman BT, Hao M, Qiu J, Jiao X, Baseler MW, Lane HC, et al. DAVID: a web server for functional enrichment analysis and functional annotation of gene lists (2021 update). Nucleic Acids Res. 2022 Jul 5;50(W1):W216–21.

44. Yuan V, Hui D, Yin Y, Peñaherrera MS, Beristain AG, Robinson WP. Cell-specific characterization of the placental methylome. BMC Genomics. 2021 Dec;22(1):6.

45. Xu RH, Chen X, Li DS, Li R, Addicks GC, Glennon C, et al. BMP4 initiates human embryonic stem cell differentiation to trophoblast. Nat Biotechnol. 2002 Dec;20(12):1261–4.

46. Shi D, Zhou X, Cai L, Wei X, Zhang L, Sun Q, et al. Placental DNA methylation analysis of selective fetal growth restriction in monochorionic twins reveals aberrant methylated CYP11A1 gene for fetal growth restriction. FASEB J. 2023 Oct;37(10):e23207.

47. Norton C, Clarke D, Holmstrom J, Stirland I, Reynolds PR, Jenkins TG, et al. Altered Epigenetic Profiles in the Placenta of Preeclamptic and Intrauterine Growth Restriction Patients. Cells. 2023 Apr 11;12(8):1130.

48. Lee S, Kim YN, Im D, Cho SH, Kim J, Kim JH, et al. DNA Methylation and gene expression patterns are widely altered in fetal growth restriction and associated with FGR development. Anim Cells Syst. 2021 May 4;25(3):128–35.

49. Monteagudo-Sánchez A, Sánchez-Delgado M, Mora JRH, Santamaría NT, Gratacós E, Esteller M, et al. Differences in expression rather than methylation at placenta-specific imprinted loci is associated with intrauterine growth restriction. Clin Epigenetics. 2019 Dec;11(1):35.

50. Roifman M, Choufani S, Turinsky AL, Drewlo S, Keating S, Brudno M, et al. Genome-wide placental DNA methylation analysis of severely growth-discordant monochorionic twins reveals novel epigenetic targets for intrauterine growth restriction. Clin Epigenetics. 2016 Dec;8(1):70.

51. Kantake M, Ikeda N, Nakaoka H, Ohkawa N, Tanaka T, Miyabayashi K, et al. IGF1 gene is epigenetically activated in preterm infants with intrauterine growth restriction. Clin Epigenetics. 2020 Dec;12(1):108.

52. Richter AE, Bekkering-Bauer I, Verkaik-Schakel RN, Leeuwerke M, Tanis JC, Bilardo CM, et al. Altered neurodevelopmental DNA methylation status after fetal growth restriction with brain-sparing. J Dev Orig Health Dis. 2022 Jun;13(3):378–89.

53. Roberts RM, Ezashi T, Temple J, Owen JR, Soncin F, Parast MM. The role of BMP4 signaling in trophoblast emergence from pluripotency. Cell Mol Life Sci. 2022 Aug;79(8):447.

54. Papadatou I, Marinakis N, Botsa E, Tzanoudaki M, Kanariou M, Orfanou I, et al. Case Report: A Novel Synonymous ARPC1B Gene Mutation Causes a Syndrome of Combined Immunodeficiency, Asthma, and Allergy With Significant Intrafamilial Clinical Heterogeneity. Front Immunol. 2021 Feb 19;12:634313.

55. Du Q, Zhang M, Gao A, He T, Guo M. Epigenetic silencing ZSCAN23 promotes pancreatic cancer growth by activating Wnt signaling. Cancer Biol Ther. 2024 Dec 31;25(1):2302924.

56. Batsukh T, Schulz Y, Wolf S, Rabe TI, Oellerich T, Urlaub H, et al. Identification and Characterization of FAM124B as a Novel Component of a CHD7 and CHD8 Containing Complex. Englert C, editor. PLoS ONE. 2012 Dec 21;7(12):e52640.

57. Care4Rare Canada Consortium, Choquet K, Tétreault M, Yang S, La Piana R, Dicaire MJ, et al. SPG7 mutations explain a significant proportion of French Canadian spastic ataxia cases. Eur J Hum Genet. 2016 Jul;24(7):1016–21.

58. Ismail MR, Ordi J, Menendez C, Ventura PJ, Aponte JJ, Kahigwa E, et al. Placental pathology in malaria: A histological, immunohistochemical, and quantitative study. Hum Pathol. 2000 Jan;31(1):85–93.

