## Supplementary figures and images for "Placental Malaria Induces a Unique Methylation Profile Associated with Fetal Growth Restriction"

### Supplemental Figure 1

##### Supplemental Figure 1.

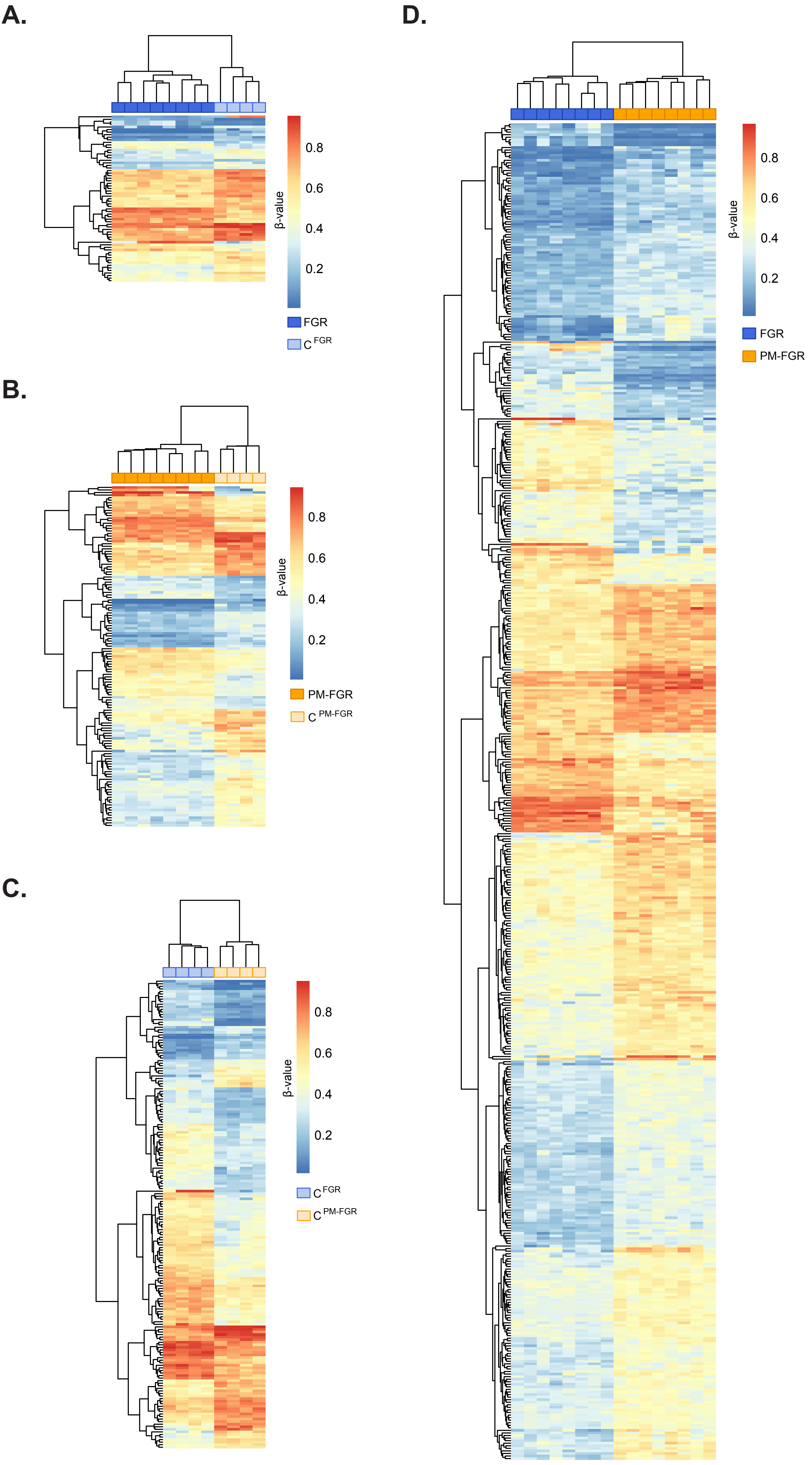

### Supplemental Figure 2

#### Supplemental Figure 2.

## A.

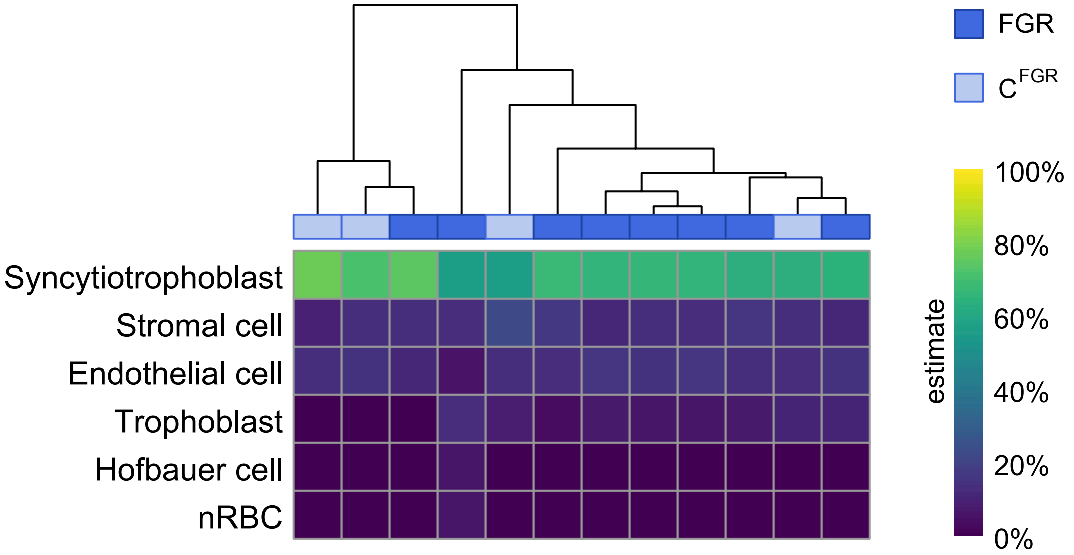


## B.

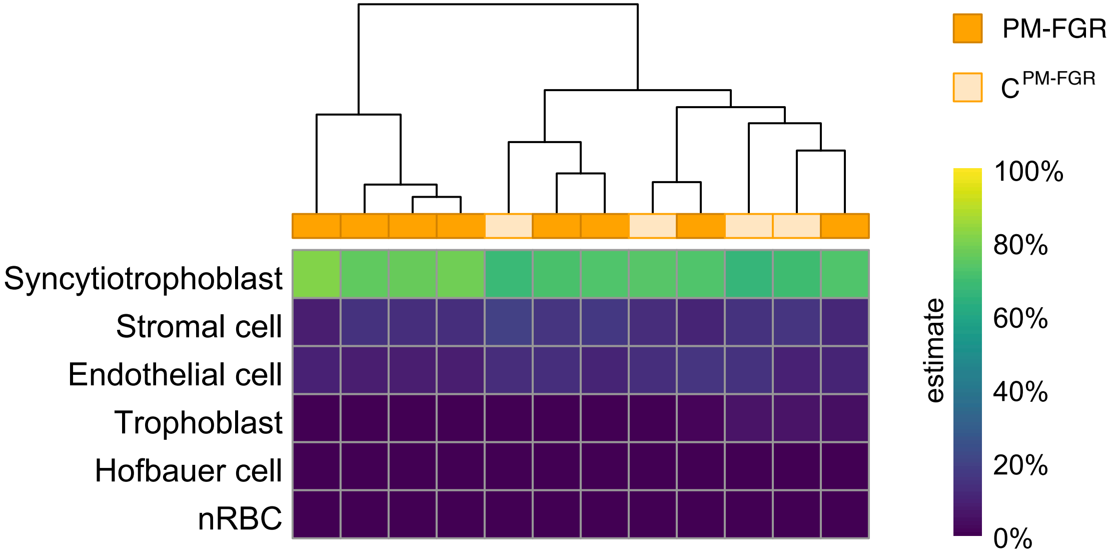
