## Supplemental Table 1 for "Placental Malaria Induces a Unique Methylation Profile Associated with Fetal Growth Restriction"

| Supplementary Table 1. Patient characteristics. | | | | | | | |
|---|---|---|---|---|---|---|---|
| Study ID | Study group | Collection site | Maternal comorbidities | Gestational week | Infant sex | Birth weight (gram) | Intergrwoth-21 birth weight percentile |
| 1 | $C^{FGR}$ | United States | anemia, generalized anxiety | 39 | Female | 3505 | 79.53 |
| 2 | $C^{FGR}$ | United States | none | 39 | Male | 2895 | 13.66 |
| 3 | $C^{FGR}$ | United States | none | 39 | Male | 3625 | 80.66 |
| 4 | $C^{FGR}$ | United States | none | 39 | Male | 3815 | 90.09 |
| 5 | $C^{PM-FGR}$ | Uganda | none | 39 | Female | 2900 | 25.41 |
| 6 | $C^{PM-FGR}$ | Uganda | none | 39 | Female | 2880 | 20.61 |
| 7 | $C^{PM-FGR}$ | Uganda | none | 38 | Female | 3650 | 90.30 |
| 8 | $C^{PM-FGR}$ | Uganda | none | 38 | Male | 2930 | 33.42 |
| 9 | FGR | United States | cHTN, SIPE, gDM, subclinical hypothyroidism | 36 | Female | 2165 | 6.36 |
| 10 | FGR | United States | paroxysmal positional vertigo | 38 | Female | 2280 | 1.62 |
| 11 | FGR | United States | none | 37 | Female | 2220 | 3.21 |
| 12 | FGR | United States | asthma | 37 | Female | 2380 | 6.01 |
| 13 | FGR | United States | gHTN, gestational thrombocytopenia | 37 | Female | 2135 | 2.59 |
| 14 | FGR | United States | Crohn’s disease, opioid use, PreE | 38 | Male | 2425 | 3.98 |
| 15 | FGR | United States | gDM, PreE, anemia | 36 | Male | 1430 | 0.14 |
| 16 | FGR | United States | gHTN, subclinical hypothyroidism, generalized anxiety, depression | 37 | Male | 1960 | 0.98 |
| 17 | PM-FGR | Uganda | anemia | 42 | Female | 2900 | 6.25 |
| 18 | PM-FGR | Uganda | none | 39 | Female | 2540 | 3.14 |
| 19 | PM-FGR | Uganda | none | 39 | Female | 2500 | 4.15 |
| 20 | PM-FGR | Uganda | none | 37 | Female | 2030 | 1.85 |
| 21 | PM-FGR | Uganda | anemia | 41 | Male | 3000 | 7.00 |
| 22 | PM-FGR | Uganda | anemia | 39 | Male | 2550 | 3.71 |
| 23 | PM-FGR | Uganda | none | 40 | Male | 2670 | 3.27 |
| 24 | PM-FGR | Uganda | anemia | 39 | Male | 2650 | 4.01 |
